# Saponin Chemistry Controls Anionic Lipid Tolerance and Divalent Metal Ion Responses in Magnetically Alignable Bicelles

**DOI:** 10.64898/2026.04.25.720808

**Authors:** Samuel D. McCalpin, Cletus Obi, Ayyalusamy Ramamoorthy

**Author notes:** **Corresponding Author:** Ayyalusamy Ramamoorthy; *E.mail:.

## Abstract

Saponin-phospholipid bicelles have emerged as promising membrane-mimetic systems for anisotropy-based NMR studies, but their utility depends on their ability to accommodate physiologically relevant lipid compositions and ionic environments. Here, we systematically investigated how saponin chemistry governs three key properties of magnetically alignable bicelles: tolerance to anionic lipid incorporation, responsiveness to lanthanide-induced alignment reorientation, and stability in the presence of divalent metal ions. Using ^31^P NMR spectroscopy as a sensitive probe of phase behavior and alignment, we compared bicelles formed with glycyrrhizic acid (GA), hederacoside C (HC), and crude Quillaja saponins (CQS). While all saponins effectively solubilized the anionic lipid DMPG, only HC supported magnetically aligned bicelles with high anionic lipid fractions (up to 70%), whereas GA was limited to low incorporation (10%). Both GA- and HC-based bicelles underwent lanthanide-induced alignment flipping, though with significant spectral broadening and intermediate alignment states indicative of increased heterogeneity. Divalent cation effects were strongly ion- and saponin-dependent; HC bicelles were robust to the presence of Ca^2+^ but were destabilized by Mg^2+^, while GA bicelles were disrupted by both ions at low concentrations. Together, these results demonstrate that saponin identity critically determines bicelle compatibility with charged lipids and ionic conditions, establishing design principles for tailoring saponin-based bicelles as versatile, biomimetic alignment media for membrane-protein structural studies.

## Introduction

Saponins are a structurally diverse class of plant-derived glycoside natural products characterized by a hydrophobic triterpenoid or steroidal aglycone group linked to one or more hydrophilic oligosaccharide chains^1–9^. This amphiphilic architecture enables strong interactions with lipid membranes, ranging from adsorption and permeabilization to full solubilization, which underlie their widespread roles in plants as antifungals and deterrents against predation.^2,10,11^ Many saponins also exhibit potent immunomodulatory properties in vertebrates and have consequently been developed as adjuvants in human and veterinary vaccines.^12–18^ For example, vaccines approved for shingles and COVID-19 include saponins derived from *Quillaja Saponaria*^16^ in their adjuvant systems.^13,19–22^

More recently, saponins have emerged as potential soft-matter building blocks for membrane-mimetic model systems. Several binary mixtures of saponins and phospholipids have been shown to self-assemble into discoidal planar bilayers, often called bicelles or nanodiscs, whose morphology resembles a planar lipid bilayer “patch” stabilized by amphiphilic “belt” molecules, such as polymers or short-chain detergents, around the rim.^6,7,23–26^ Such assemblies are attractive for membrane biophysics because they provide a hydrated bilayer environment while remaining experimentally tractable across a wide range of analytical methods.^6,7,23^ In particular, magnetically alignable bicelles enable anisotropy-based NMR experiments that exploit partial orientational order to measure residual dipolar couplings (RDCs), residual quadrupolar couplings, and other orientation-dependent interactions.^6,27–29^

To date, the development of saponin-phospholipid bicelles has largely emphasized zwitterionic phosphocholines such as 1,2-dimyristoyl-sn-glycero-3-phosphocholine (DMPC). In our previous work, we demonstrated the ability of crude Quillaja saponin (CQS) extracts to promote the formation of a magnetically aligning bicellar phase that enabled the measurement of RDCs for the water-soluble protein cytochrome C.^6^ We also found that using the pure triterpenoid saponins aescin, glycyrrhizic acid (GA), ginsenoside Rb1 (Grb1), or hederacoside C (HC) in place of CQS produces a more homogeneous alignable bicellar phase.^7^ Complementary studies by Hellweg and co-workers have further characterized the aescin-DMPC and GA-1,2-dioleoyl-sn-glycero-3-phosphoglycerol (DOPG)^30^ system, including bicelle formation and bicelle-vesicle interconversion as a function of temperature, composition, and dilution.^23,25^ Collectively, these findings suggest that saponins can serve as versatile alternatives to conventional detergent-like rim-stabilizers, potentially expanding the compositional and functional space of bicelle-based model membranes. However, to establish the breadth of their utility for physiologically relevant membrane protein studies and anisotropy-based NMR applications, several key properties of saponin-phospholipid bicelles must be determined.

First, the bicelles must allow incorporation of anionic lipids. Biological membranes possess a net negative surface charge arising from substantial fractions of anionic lipids, including phosphatidylglycerols, phosphatidylserines, phosphatidylinositols, phosphatidic acids, and acidic glycolipids, whose distribution regulates signaling, protein recruitment, and membrane electrostatics.^31–34^ Model membrane systems intended to emulate physiological conditions must therefore tolerate, and ideally allow systematic tuning of, negative lipid content. However, the formation of well-aligned bicelle phases at elevated fractions of anionic lipids is not guaranteed.^31,35–37^ In some systems, strong electrostatic repulsions or altered packing constraints due to increased anionic lipid content can shift the phase behavior away from bicelles and toward alternative morphologies, including vesicles, micelles, or heterogeneous mixtures. In saponin-containing mixtures, electrostatic interactions between acidic functional groups in the saponin molecules in the belt and anionic lipid headgroups in the bilayer surface may reduce lipid solubilization and disfavor bicelle formation.^38^ Indeed, Gräbitz-Bräuer et al. reported that the saponin β-aescin, which forms homogeneous bicelles with DMPC, did not mix at all with the anionic lipid DOPG.^38^ Similarly, DHPC was unable to form bicelles with pure DMPG or DMPS but could form bicelles with lipid mixes comprised of 75% DMPC and 25% DMPG or DMPS.^37,39^ Therefore, it is important to determine the anionic lipid content that is tolerated in bicelles formed by saponins of interest.

Second, it is advantageous to be able to control the orientation of bicelle alignment in the magnetic field. In the absence of additives, phospholipid acyl chains possess a negative magnetic susceptibility anisotropy (Δχ), which drives bilayer assemblies to align with their membrane normal perpendicular to the external magnetic field (B_0_). This alignment produces a weakly ordered environment in which solutes experience residual anisotropic interactions, enabling measurement of RDCs and related observables. Importantly, the magnitude of these splittings depends on the angular factor (3cos^2^θ − 1)/2, meaning that molecules undergoing fast axially symmetric motion relative to the bilayer normal (such as small molecules or peptides) will see their anisotropic splittings reduced by a factor of −1/2. Some membrane proteins with transmembrane helices possess a positive Δχ that cancels out the negative anisotropy of the lipids, disrupting the magnetic alignment of the system.^40^ To combat these issues, it has been found that doping bicelles with lanthanide ions, which have positive Δχ, flips their alignment by 90°, such that the bilayer normal aligns parallel with B_0_. In this orientation the (3cos2θ - 1)/2 scaling factor becomes 1, and transmembrane helices cannot disrupt alignment due to their positive Δχ. Several lanthanides, including Yb^3+^, Tm^3+^, Eu^3+^, and others, have been reported to induce such reorientation in classical phospholipid bicelles.^41–43^ Yet, compatibility is not universal: some belt molecules and additives can bind lanthanides strongly enough to sequester them, preventing the flip or producing heterogeneous alignment behavior. For example, we previously observed that CQS-DMPC bicelles doped with Yb^3+^ would only exhibit a flipped orientation after a filtration step to remove unbound CQS.^6^ Because saponins contain a diversity of oxygen-rich functional groups and can vary substantially in charge and chelation propensity, it is important to establish whether saponin-stabilized bicelles undergo the expected lanthanide-induced reorientation for each system.

Finally, divalent metal ion tolerance is a practical requirement for many applications of bicelle and nanodisc systems.^44–47^ Many membrane proteins require millimolar concentrations of divalent cations such as Ca^2+^ and Mg^2+^ for folding, ligand binding, catalysis, or regulation, and these ions can also influence membrane electrostatics and headgroup interactions. In classical DHPC phospholipid bicelles, divalent cations have been reported to modulate bicelle morphology and magnetic alignment, with effects that depend on both ion identity and concentration^48–51^. Sensitivity to divalent metal ions is also a well-recognized limitation for several nanodisc scaffolds.^52^ For example, commonly used styrene-maleic acid (SMA)-based polymer nanodiscs can destabilize or precipitate in the presence of Ca^2+^ or Mg^2+^ due to ion-mediated charge neutralization and polymer aggregation.^52,53^ Because saponins contain multiple oxygen-rich functional groups and may present charged or chelating motifs depending on structure, establishing whether saponin-phospholipid bicelles remain intact under divalent-cation conditions is essential for defining their uses in membrane-protein biophysics and NMR applications.

In this work, we extend the characterization of saponin-phospholipid bicelles by systematically evaluating three properties that are essential for their use as physiologically relevant, magnetically alignable model membranes: (i) tolerance to incorporation of anionic lipids, (ii) the ability to undergo lanthanide-induced reorientation in a magnetic field, and (iii) stability in the presence of divalent metal ions. We use ^31^P NMR spectroscopy as the primary reporter of phase state and magnetic alignment, since phospholipid headgroup’s ^31^P chemical shift anisotropy provides a sensitive reporter of bicelle formation, bilayer orientation, and sample heterogeneity. To probe how saponin chemistry governs these behaviors, we focus on two structurally distinct triterpenoid saponins that highlight key electrostatic and practical considerations: glycyrrhizic acid (GA), an inexpensive but acidic saponin expected to accentuate electrostatic interactions with anionic lipid headgroups and metal ions, and hederacoside C (HC), a comparatively costly but effectively nonionic saponin expected to minimize charge-driven effects. In addition, crude Quillaja saponin (CQS) was considered as a low-cost, predominantly nonionic reference system that forms magnetically alignable but less homogeneous bicellar phases. Together, these comparisons establish how saponin charge and purity influence bicelle compatibility with anionic membranes, orientation control via lanthanide doping, and robustness to divalent cation conditions relevant to membrane-protein function.

## Experimental Methods

### Materials

The phospholipids 1,2-dimyristoyl-sn-glycero-3-phosphocholine (DMPC) and 1,2-dimyristoyl-sn-glycero-3-phospho-(1’-rac-glycerol) (DMPG) were purchased from Avanti Polar Lipids (Alabaster, AL). Glycyrrhizic acid monoammonium salt was obtained from Thermo Fisher Scientific (Waltham, MA), while Quillaja saponins and hederacoside C were sourced from MilliporeSigma (Burlington, MA). All materials were used as received, without further purification.

### Transmittance Assay

Phospholipids were obtained as lyophilized powders, weighed out, and stocks were prepared in sodium phosphate buffer (10 mM NaPO_4_, 100 mM NaCl, pH 7.4) for a lipid concentration of 10 mg/mL as reported previously.^7^ The lipid stock was homogenized by three freeze-thaw cycles using liquid N_2_ and a warm water bath. Saponin powders were measured out and dissolved directly in the lipid stock for a saponin concentration of 50 mg/mL. This solution was then subjected to three freeze-thaw cycles. Then, samples for the transmittance assay were prepared by diluting the saponin solution with the lipid stock. These samples were further mixed in a thermal mixer at 1000 rpm and 30 °C (or at least 5 °C above the main lipid phase transition temperature) for at least 30 min. Samples were plated in triplicate in a non-binding, clear-bottomed, black-walled 384-well microplate. Absorbance was read at 790 nm using a BioTek Synergy 2 microplate reader. Temperature was set to 25 °C during the absorbance measurement. Buffer was used as a control and its absorbance subtracted from the sample absorbance. Two absorbance measurements were performed for each sample and averaged. Transmittance was calculated and normalized from the absorbance according to:

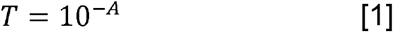

### ^*31*^*P NMR*

NMR samples were prepared from phospholipid stocks in tris buffer (10 mM tris, 100 mM NaCl, pH 7.4) that were homogenized by three freeze-thaw cycles. Lanthanide salts were added to the buffer in the relevant samples. The phospholipid stocks were used to dissolve saponin powder to achieve a final phospholipid concentration of 100 mg/mL and the appropriate ratio of saponin. Saponin-lipid mixtures were subjected to three additional freeze-thaw cycles. Samples were packed in a 2.5 mm thin-walled Bruker rotor (approximately 25 µL sample volume, containing 2.5 mg phospholipid). ^31^P NMR spectra were acquired on a 400 MHz Bruker spectrometer using a triple-resonance HXY 2.5 mm Bruker MAS probe under static conditions. Spectra were collected with a 2.2 µs 90° pulse, 3 s recycle delay, 1024 scans, and 35 kHz ^1^H decoupling. An exponential window function with 20 Hz line broadening was used to process the data.

## Results and Discussion

### Incorporation of anionic lipids into saponin-phospholipid bicelles

In our previous study using CQS and the anionic lipid DMPG, we observed a strongly non-monotonic solubilization profile in aqueous buffer: DMPG became increasingly soluble as the CQS fraction was increased up to ∼40% (relative to total lipid + saponin), was fully solubilized near ∼40-50% CQS, then became progressively less soluble at higher CQS fractions (∼50-80%), with little to no solubilization observed above ∼80% CQS. These results suggested that electrostatic interactions and/or changes in aggregate morphology impose constraints on how acidic saponin mixtures accommodate anionic headgroups and raised the question of whether alternative saponins could more robustly support anionic lipid incorporation while preserving bicelle alignment.

We first examined the ability of GA and HC (Figure 1a) to solubilize DMPG. Both saponins dissolved DMPG efficiently, and the concentration requirements closely mirrored those previously observed for solubilization of the zwitterionic lipid DMPC (Figure 1b). This result indicates that, at the level of bulk lipid dispersion and solubilization, neither GA nor HC is intrinsically incompatible with an anionic phosphatidylglycerol headgroup. However, solubilization alone is not sufficient to establish the formation of magnetically alignable planar bilayers. Consistent with this distinction, ^31^P NMR spectra of binary saponin-DMPG mixtures did not exhibit the characteristic sharp anisotropic resonances associated with a magnetically aligned bicellar phase for any of the CQS, GA, or HC (Figures 1c and S1), even when using the same saponin:lipid ratios that previously yielded the most homogeneous aligned phases in the corresponding saponin-DMPC systems. Moreover, systematic variation of the CQS:DMPG ratio likewise failed to produce an aligned phase (Figure S2), indicating that the absence of alignment is not simply due to an off-optimal mixing ratio for this lipid.

**Figure 1.**
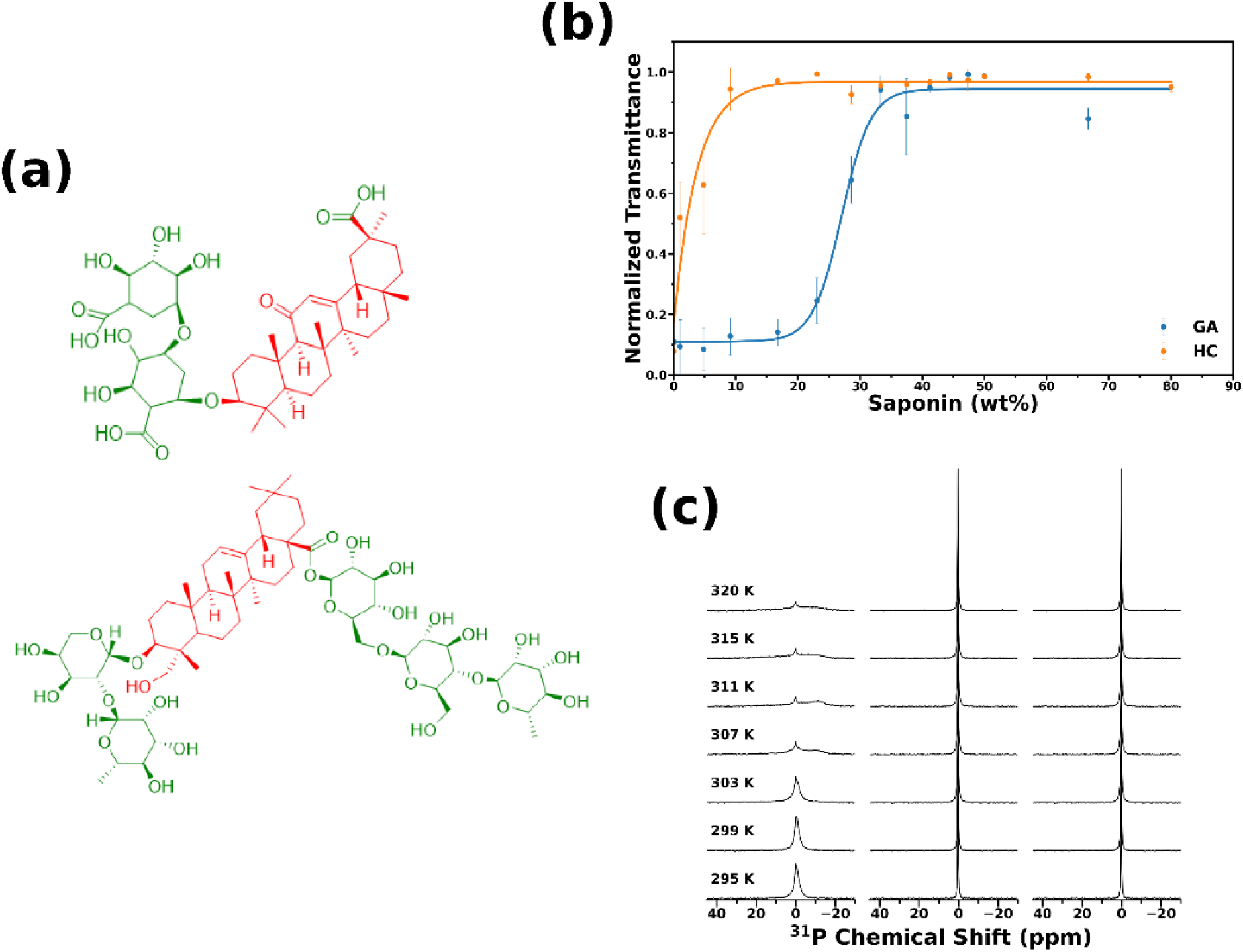
Solubilization of DMPG by Saponins. (a) Molecular structures of GA (top) and HC (bottom). Hydrophobic portions are colored in red, hydrophilic in green. (b) Normalized transmittance measured at 720 nm of 10 mg/mL DMPG mixed with varied concentrations of GA (blue) and HC (orange). (c) Temperature-dependent ^31^P NMR spectra of 100 mg/mL DMPG plus 50 mg/mL CQS (left), 20 mg/mL GA (center), or 20 mg/mL HC (right). All spectra were collected with a 400 MHz spectrometer. Spectra collected with additional temperature points are presented in Figure S1.

We next tested whether aligned phases could be supported in *ternary* mixtures containing both DMPC and DMPG while maintaining the same saponin:total phospholipid ratios used in the optimized DMPC bicelles (Figures 2 and 3). For mixtures with GA (Figure 2), an aligned phase formed with mixtures of DMPC and DMPG, but only within a narrow composition window. Specifically, mixtures containing 10% DMPG (relative to total DMPC + DMPG) displayed distinct anisotropic peaks in ^31^P NMR spectra corresponding to aligned DMPC and DMPG headgroups (approximately −10.9 ppm and −9.5 ppm, respectively) across the temperature range 303 K – 320 K. In contrast, no aligned phase was detected at higher DMPG fractions, suggesting that GA-stabilized bicelles tolerate only limited incorporation of anionic lipid before the aligned planar morphology is lost or replaced by a more isotropic and/or heterogeneous mixture of phases.

**Figure 2.**
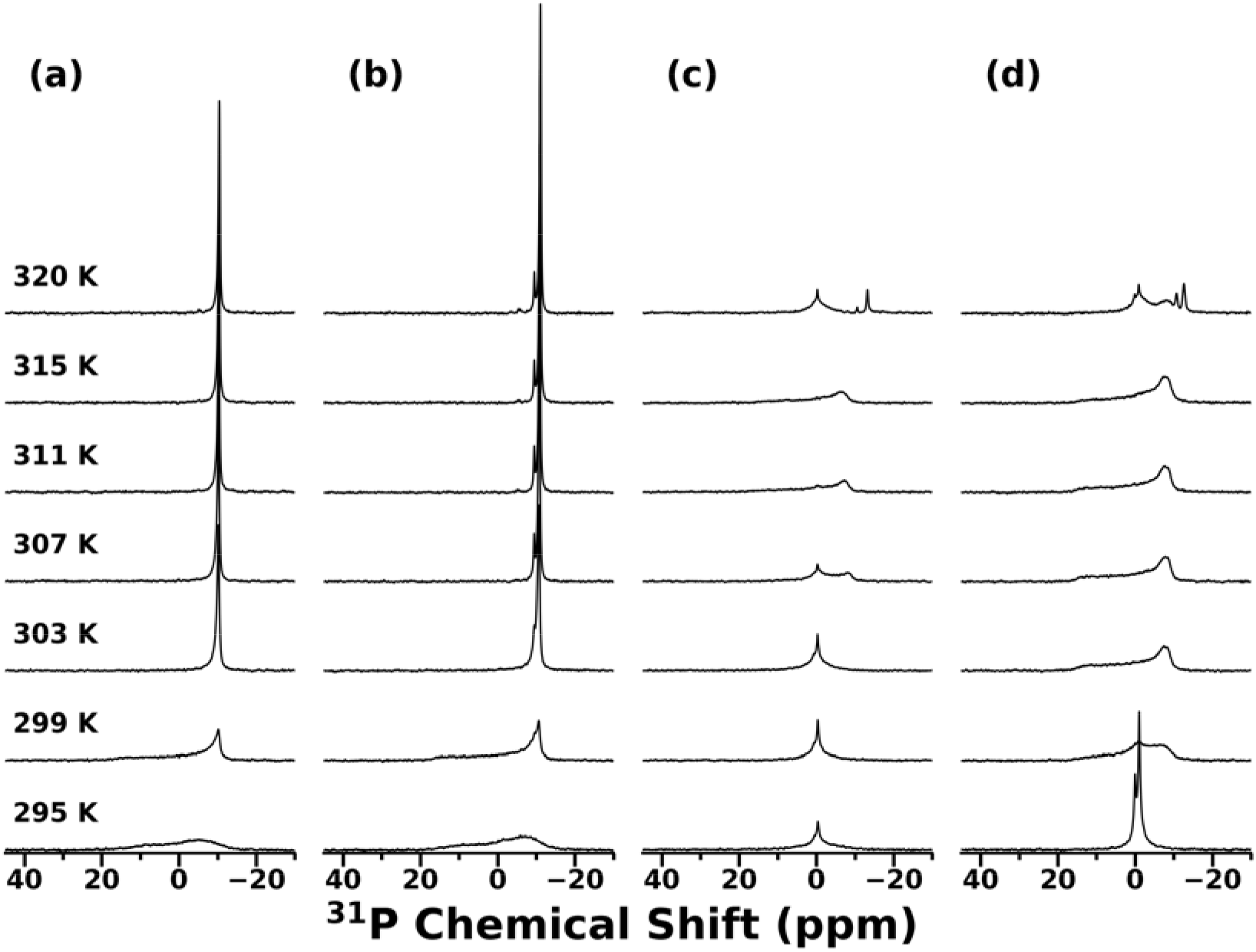
Effect of DMPG on bicelle formation by mixtures of DMPC and GA. Temperature-dependent ^31^P NMR spectra were collected with a 400 MHz spectrometer for samples containing 20 mg/mL GA and 100 mg/mL total phospholipid. The phospholipid comprised DMPC and (a) 0%, (b) 10%), (c) 15%, or (d) 30% DMPG. Spectra collected with additional temperature points are presented in Figure S3.

**Figure 3.**
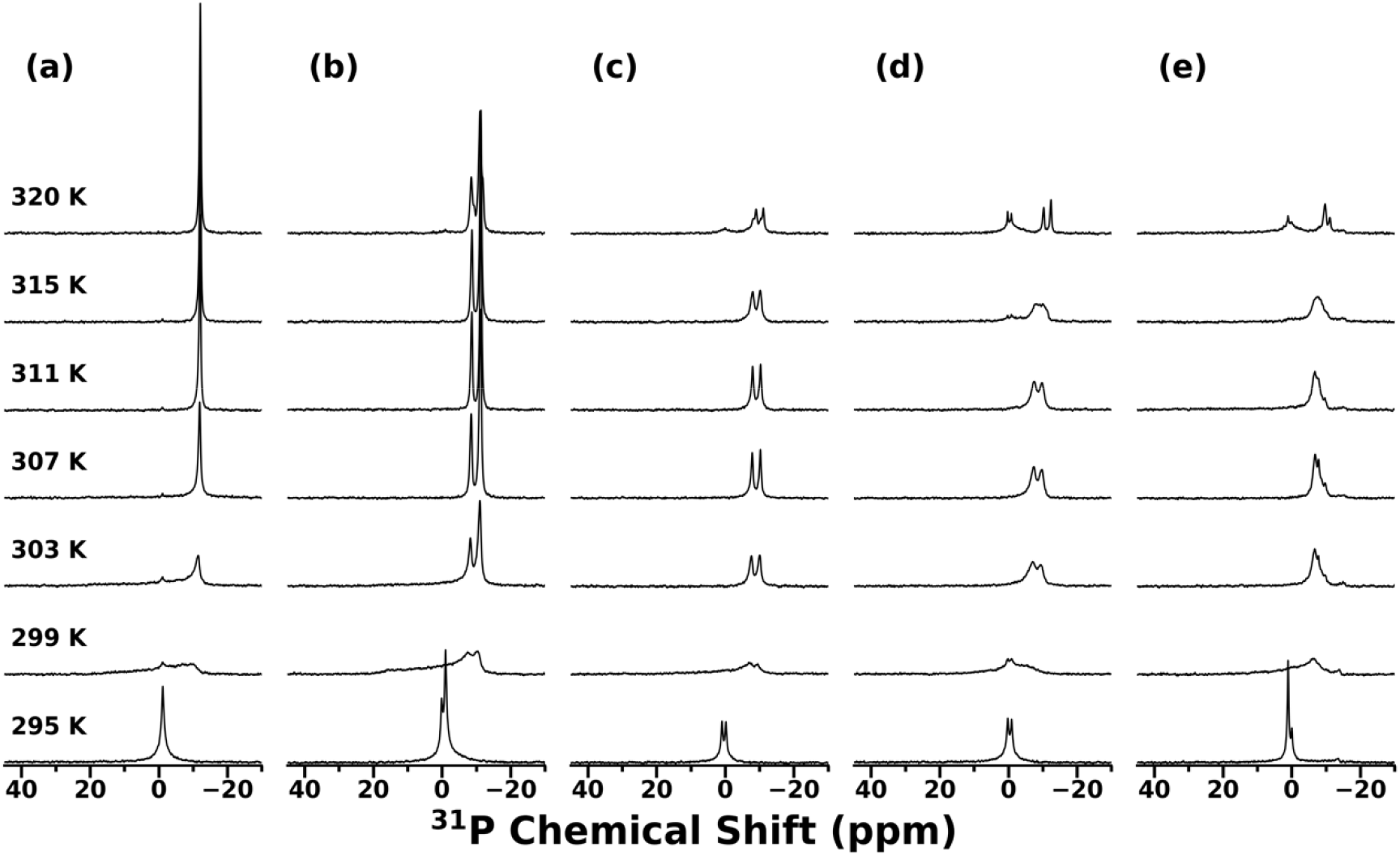
Effect of DMPG on bicelle formation by mixtures of DMPC and HC. Temperature-dependent ^31^P NMR spectra were collected with a 400 MHz spectrometer for samples containing 20 mg/mL HC and 100 mg/mL total phospholipid. The phospholipid comprised DMPC and (a) 0%, (b) 30%), (c) 50%, (d) 55%, or (e) 70% DMPG. Spectra collected with additional temperature points are presented in Figure S4.

In contrast, mixtures with HC exhibited substantially greater tolerance to anionic lipid incorporation (Figure 3). Well-resolved anisotropic peaks consistent with magnetic alignment were observed across compositions with up to 70% DMPG. For example, at 30% DMPG, distinct aligned resonances for DMPC and DMPG were observed near - 11.1 ppm and −8.5 ppm, respectively, across 303 K – 320 K. As the DMPG fraction increased, the aligned peaks progressively broadened and shifted to higher frequency. At the highest DMPG content tested (70%), the DMPC and DMPG resonances became substantially overlapped, converging around −7.4 ppm. This observation is consistent with increased DMPG content and surface charge density causing greater headgroup disorder, altered lipid environments (e.g. DMPC and DMPG separating into distinct domains), and/or a gradual change in bicelle morphology and surface dynamics. Nevertheless, the persistence of anisotropic spectral signatures over this wide composition range demonstrates that HC can support magnetically alignable bicellar phases even at high anionic lipid fractions, whereas GA is restricted to only low levels of DMPG incorporation.

Taken together, these data show that although GA and HC both solubilize DMPG effectively, the ability to form aligned planar bilayers in the presence of anionic lipid depends strongly on saponin identity. The markedly higher anionic-lipid tolerance of HC relative to GA is consistent with a model in which minimizing charge-driven incompatibilities between belt molecules and anionic headgroups is critical for maintaining a stable bicellar phase as membrane surface charge density increases.

### Lanthanide-induced orientation flipping of saponin-DMPC bicelles

In our prior work on CQS-DMPC bicelles, addition of Yb^3+^ produced a clear orientation flip that was readily monitored by ^31^P NMR: the characteristic aligned resonance shifted from approximately −14 ppm in the perpendicular state to approximately +15 ppm in the flipped, parallel-aligned state, with the extent of flipping depending on the concentration of added Yb^3+^. Notably, however, this reorientation was only observed after a filtration step to remove excess CQS that was not bound to bicelles, suggesting that free saponin can sequester lanthanide ions and suppress the effective lanthanide-bicelle interaction needed to drive the flip.

Here we evaluated lanthanide-induced reorientation in the more compositionally defined GA-DMPC and HC-DMPC bicelle systems using thulium(III) (Tm^3+^), which was selected because it provides a particularly large positive Δχ and is therefore expected to be an efficient inducer of alignment flipping. ^31^P NMR spectra of GA-DMPC bicelles showed a single sharp anisotropic peak near −11 ppm in the absence of lanthanide (Figure 4a), consistent with a homogeneous, perpendicular-aligned bicelle phase under these conditions. Upon addition of 3 mM Tm^3+^, this resonance shifted to approximately +5 ppm (Figure 4c), indicating formation of a predominantly flipped, parallel-aligned state. Compared to the lanthanide-free sample, the flipped resonance was substantially broadened. At an intermediate lanthanide concentration (2 mM Tm^3+^), GA-DMPC exhibited mixed alignment behavior, with both the original perpendicular peak and the flipped peak simultaneously present (Figure 4b). This coexistence implies that the flipping transition occurs over a finite concentration window and is not strictly cooperative under these conditions, consistent with heterogeneous lanthanide binding and/or gradual evolution of bicelle morphology as Tm^3+^ is introduced.

**Figure 4.**
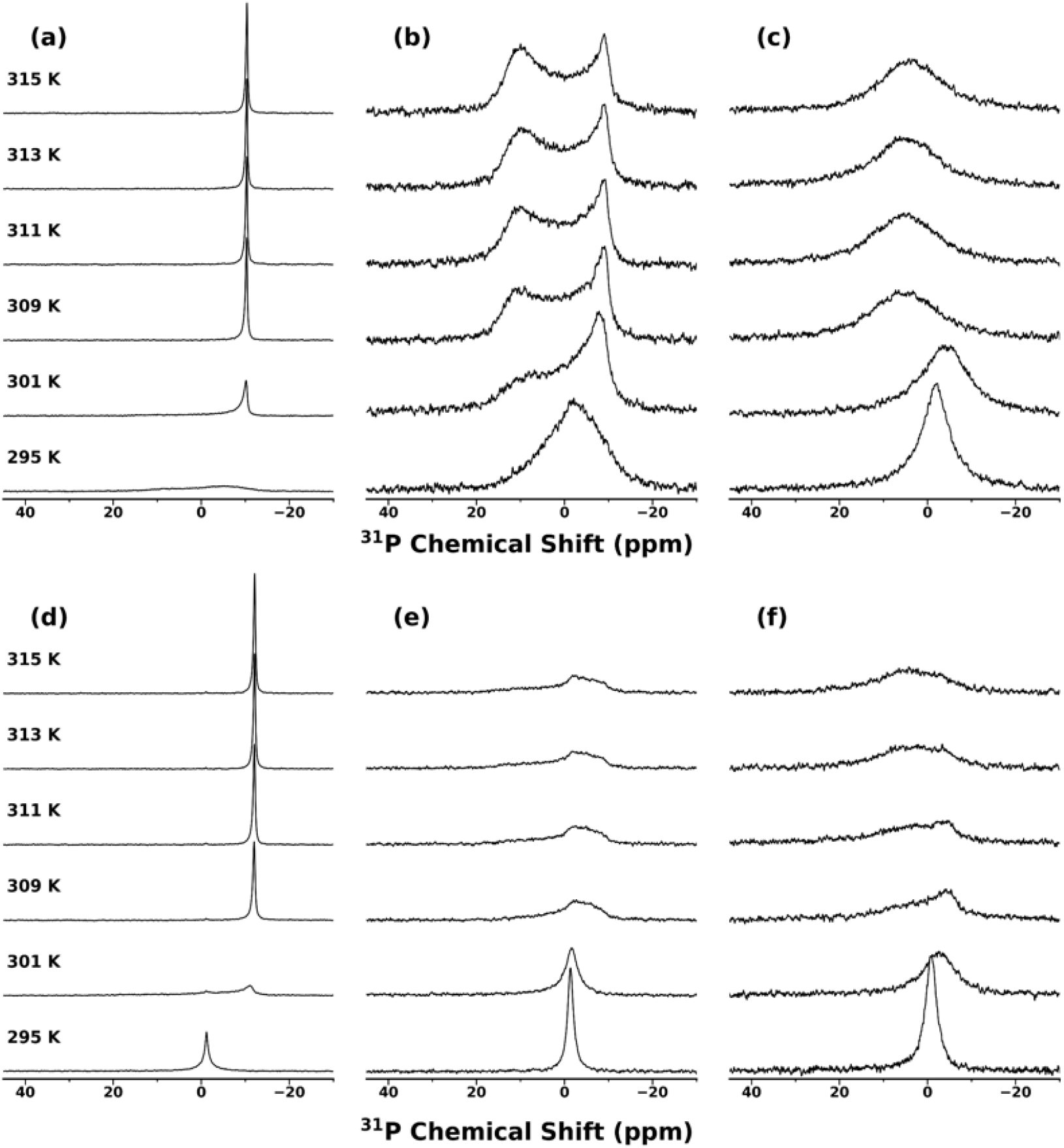
Orientation flipping of saponin bicelles. Temperature-dependent ^31^P NMR spectra were collected with a 400 MHz spectrometer for samples containing 100 mg/mL DMPC and (a) 20 mg/mL GA and 0 mM TmCl_3_, (b) 20 mg/mL GA and 2 mM TmCl_3_, (c) 20 mg/mL GA and 3 mM TmCl_3_, (d) 20 mg/mL HC and 0 mM TmCl_3_, (e) 20 mg/mL HC and 1 mM TmCl_3_, (f) 20 mg/mL HC and 2 mM TmCl_3_.

HC-DMPC bicelles displayed qualitatively similar behavior but with notable differences in the concentration dependence and spectral line shape. In the absence of Tm^3+^, HC-DMPC bicelles produced an aligned resonance near −12 ppm (Figure 4d). Upon addition of 2 mM Tm^3+^, the spectrum shifted toward a resonance near +2.5 ppm (Figure 4f), again consistent with lanthanide-induced reorientation to a parallel-aligned state. Interestingly, at 1 mM Tm^3+^, the HC-DMPC spectrum did not present as a simple superposition of two narrow aligned peaks; instead, it exhibited a powder pattern-like line shape (Figure 4e). This lineshape is indicative of a broader distribution of orientations and/or incomplete macroscopic alignment, suggesting that HC-DMPC passes through an intermediate regime in which the sample is neither uniformly perpendicular-aligned nor uniformly parallel-aligned. One plausible interpretation is that, at intermediate Tm^3+^ concentrations, partial lanthanide binding perturbs bicelle ordering and produces a mixture of aligned and less-aligned aggregates, which then resolves into a predominantly flipped state once sufficient lanthanide is present to enforce parallel orientation. Also, as observed for GA-DMPC, the flipped peak was substantially broader than the lanthanide-free aligned peak.

The pronounced linewidth broadening observed upon the addition of Tm^3+^ in both cases is consistent with both paramagnetic line broadening and with increased heterogeneity in the aligned phase, either through a distribution of bicelle sizes and orientations, altered lipid order parameters, or partial coexistence between flipped and non-flipped populations. Paramagnetic broadening is expected given the presence of Tm^3+^, but mixed orientations for GA and loss of orientation for HC observed with intermediate Tm^3+^ concentrations indicate that lanthanide doping at least partially compromises the uniformity of the aligned phase even when flipping is achieved.

Taken together, these data demonstrate that both GA-DMPC and HC-DMPC bicelles can undergo lanthanide-induced orientation flipping without requiring a filtration step to remove excess saponin, in contrast to our earlier observations with CQS-DMPC. However, in both purified saponin systems, flipping is accompanied by pronounced spectral broadening and, in some cases, intermediate powder-pattern line shapes, indicating that lanthanide doping can introduce structural heterogeneity even as it achieves the desired control over alignment orientation. These findings suggest that while Tm^3+^ provides a reliable means of switching saponin bicelles into a parallel-aligned state, optimizing lanthanide identity, concentration, and sample preparation conditions may be necessary to preserve maximal alignment homogeneity in applications requiring high spectral resolution.

### Stability of saponin bicelles against divalent metal ions

To evaluate the compatibility of saponin-phospholipid bicelles with conditions relevant to metal-dependent membrane proteins, we investigated the stability of magnetically aligned bicellar phases in the presence of increasing concentrations of divalent cations. Because divalent ions can influence membrane electrostatics, lipid packing, and the aggregation state of amphiphilic belt molecules, their presence can either subtly modulate bicelle morphology or destabilize the aligned phase entirely. We therefore used ^31^P NMR spectroscopy to monitor the persistence of alignment in HC-DMPC bicelles as a function of added Ca^2+^ (Figure 5) and Mg^2+^ (Figure 6).

**Figure 5.**
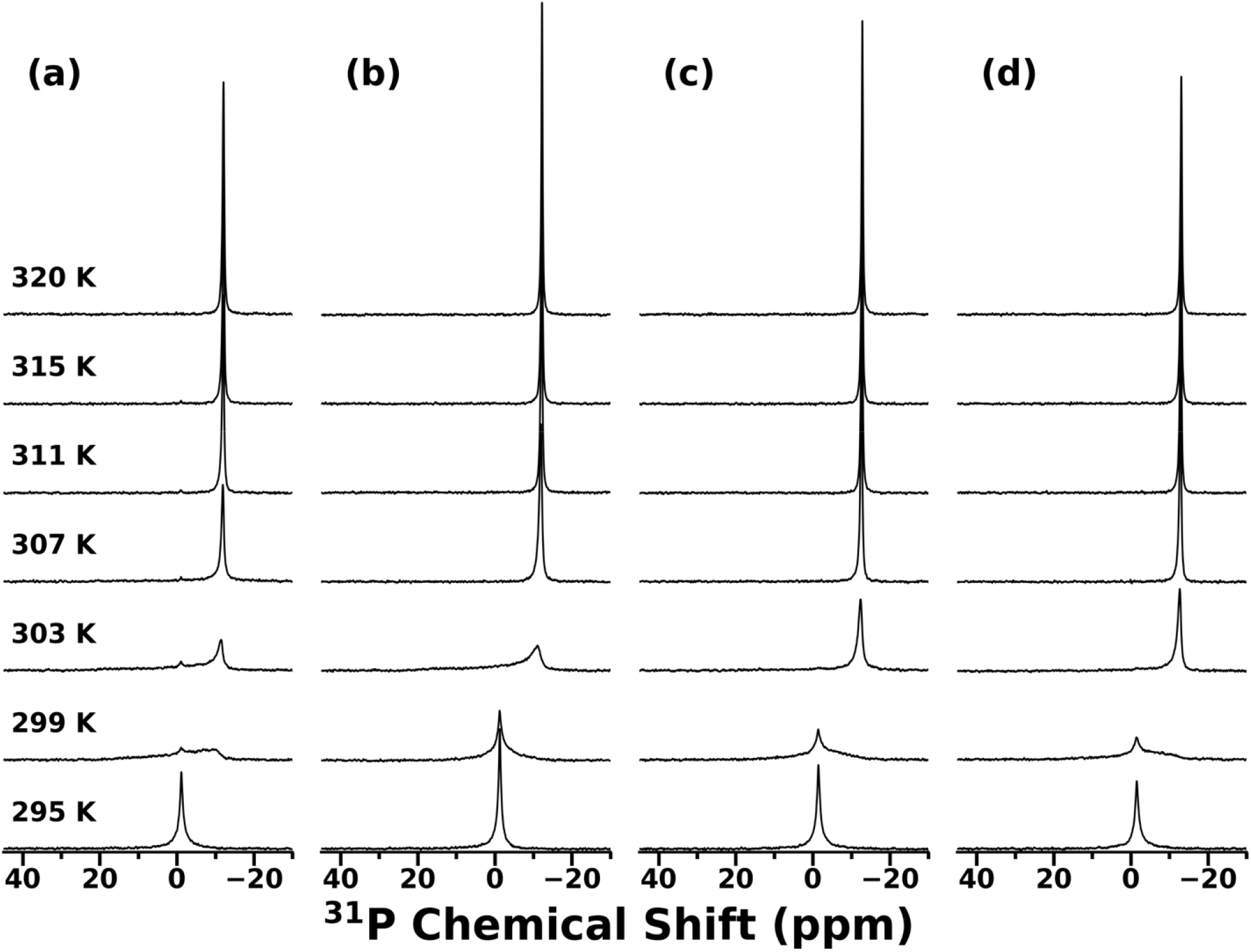
Effect of Ca^2+^ on the stability of saponin bicelles. Temperature-dependent ^31^P NMR spectra were collected for samples containing 100 mg/mL DMPC, 20 mg/mL HC, and (a) 0 mM, (b) 10 mM, (c) 50 mM, or (d) 100 mM CaCl_2_. Spectra collected with additional temperature points are presented in Figure S5.

**Figure 6.**
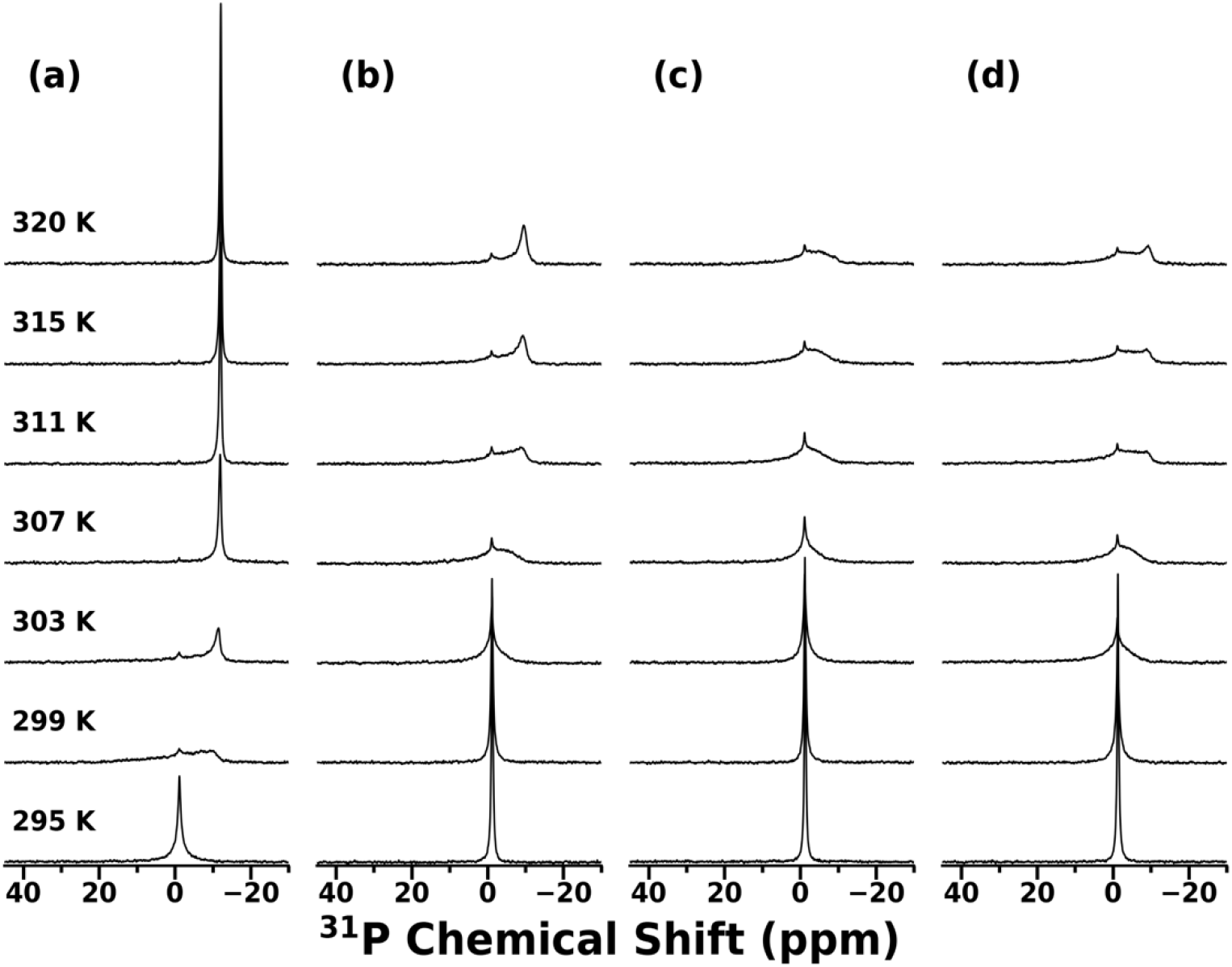
Effect of Mg^2+^ on the stability of saponin bicelles. Temperature-dependent ^31^P NMR spectra were collected for samples containing 100 mg/mL DMPC, 20 mg/mL HC, and (a) 0 mM, (b) 10 mM, (c) 50 mM, or (d) 100 mM MgCl_2_. Spectra collected with additional temperature points are presented in Figures S6 and S7.

Across the concentration range tested, Ca^2+^ had little observable impact on the ability of HC bicelles to form and maintain an aligned phase. The ^31^P spectra recorded in the absence of Ca^2+^ were qualitatively similar to those obtained in the presence of 10, 50, or 100 mM Ca^2+^ (Figure 5), with alignment retained over comparable temperature windows and no major changes to line shape indicative of vesiculation or phase disruption. The only reproducible effect of Ca^2+^ addition was a modest shift in the lower temperature boundary for alignment: at higher Ca^2+^ concentrations (50 and 100 mM), the onset of alignment occurred at slightly lower temperatures (∼303 K) than in the Ca^2+^-free sample (∼307 K). This small downward shift suggests that Ca^2+^ may weakly stabilize the aligned bicellar phase or facilitate alignment kinetics at lower temperature, but overall Ca^2+^ does not appear to compromise the structural integrity or homogeneity of HC-stabilized bicelles under the tested conditions.

In contrast to Ca^2+^, Mg^2+^ exerted a pronounced destabilizing effect on bicelle alignment (Figure 6). In Mg^2+^-free samples, HC bicelles exhibited alignment over a low-temperature window (303 K – 307 K), as evidenced by the characteristic anisotropic ^31^P resonance of an oriented bilayer assembly.

Upon the addition of Mg^2+^, this same temperature range no longer supported a well-defined aligned phase. Instead, the spectra transitioned from a predominantly isotropic resonance to a narrow, powder pattern-like line shape superimposed on a low-intensity isotropic peak. This type of intermediate lineshape is consistent with incomplete macroscopic alignment and/or coexistence of aligned and un-aligned morphologies, indicating that Mg^2+^ perturbs the ordered bicellar phase even at relatively low concentrations.

At 10 mM Mg^2+^, the aligned phase was not entirely eliminated but appeared to be displaced to higher temperature: alignment was only clearly observed above ∼320 K (Figure S7), suggesting that Mg^2+^ raises the thermal threshold required to access a stable aligned morphology. However, at Mg^2+^ concentrations exceeding 10 mM, no aligned phase was observed within the full temperature range investigated, even up to 360 K. Mechanistically, the markedly different effects of Ca^2+^ and Mg^2+^ on HC bicelles likely reflect differences in how these ions interact with the lipid headgroup region and with the saponin-stabilized rim environment. Although both ions are divalent, Mg^2+^ is more strongly hydrated and typically forms tighter inner-sphere coordination complexes with oxygen donors, which can enhance its ability to perturb interfacial hydration structure, headgroup dynamics, and electrostatic screening at the membrane surface. Such changes can reduce the uniformity of bicelle orientation and promote transitions toward less well-aligned or more heterogeneous morphologies, consistent with the emergence of powder-pattern-like lineshapes and loss of a single sharp aligned resonance at intermediate Mg^2+^ concentrations. In addition, Mg^2+^ may interact more strongly with oxygen-rich functional groups present in HC, effectively altering the balance of rim stabilization versus bilayer packing and shifting the temperature window required for a stable aligned phase. By contrast, the comparatively minor effect of Ca^2+^ - limited to a small decrease in the lower temperature bound for alignment even at high concentrations – suggests that Ca^2+^ does not substantially disrupt the HC-lipid assembly and may weakly stabilize the aligned phase by screening repulsive interactions and/or promoting more favorable lipid packing at the bicelle surface.

To determine whether electrostatic interactions with the saponin belt might disturb bicelle formation, we also performed the same set of experiments for GA-DMPC bicelles (Figures S8 and S9). Both divalent metal ions, even at the lowest concentration tested (10 mM), caused the ^31^P lineshapes to broaden or regain the left edge of a powder pattern, indicating partial disruption of bicelle formation. Together, these observations indicate a twofold effect. First, for saponins with a negative charge, all divalent metal ions significantly inhibit bicelle formation, likely by electrostatic-driven competition with the lipids for saponin binding. On the other hand, divalent-ion sensitivity in nonionic saponin bicelles is not dictated primarily by ionic charge but instead depends on ion-specific coordination and hydration properties that modulate interfacial structure and the stability of the aligned bicellar morphology.

## Conclusion

In this study, we expanded the characterization of saponin-phospholipid bicelles by evaluating three properties that govern their utility as physiologically relevant, magnetically alignable membrane mimetics: tolerance to anionic lipid incorporation, lanthanide-induced orientation flipping, and stability against divalent metal ions. Using ^31^P NMR as a sensitive reporter of phase state, bilayer orientation, and sample heterogeneity, we show that these behaviors depend strongly on saponin identity and cannot be inferred from lipid solubilization alone.

First, although binary mixtures of CQS, GA, or HC with the anionic lipid DMPG did not yield an aligned bicellar phase under conditions that produce well-aligned DMPC bicelles, incorporation of DMPG was achievable in ternary mixtures with DMPC in a saponin-dependent manner. GA-based bicelles tolerated only low anionic lipid content, exhibiting aligned resonances only at 10% DMPG, whereas HC-based bicelles maintained magnetic alignment across a wide range of compositions up to 70% DMPG. This pronounced difference highlights the importance of minimizing charge-driven incompatibilities between the rim-stabilizing belt molecules and anionic headgroups when designing bicelles with physiologically relevant surface charge.

Second, both GA-DMPC and HC-DMPC bicelles underwent lanthanide-induced alignment flipping upon the addition of Tm^3+^, demonstrating that purified saponin systems can access the parallel-aligned state without the filtration step previously required for CQS-DMPC bicelles doped with Yb^3+^. However, flipping was accompanied by substantial linewidth broadening and intermediate-regime line shapes consistent with a combination of paramagnetic effects and partial loss of alignment uniformity during the flipping transition. These observations indicate that lanthanide doping provides a viable route to control the orientation of aligned saponin bicelles, but that optimization of lanthanide identity and concentration may be necessary to preserve maximal spectral resolution for demanding NMR applications.

Finally, we found that the divalent metal ion tolerance of saponin bicelles is highly ion- and saponin-specific. HC bicelles retained alignment in the presence of Ca^2+^ concentrations up to 100 mM with only minor changes to the temperature boundary for alignment, whereas Mg^2+^ strongly destabilized the aligned phase, eliminating alignment above 10 mM and shifting the aligned regime to higher temperatures at lower concentrations. GA bicelles lost alignment homogeneity in the presence of low concentrations of both divalent metal ions.

Together, these results establish that saponin-lipid bicelles offer a tunable platform whose performance can be tailored through careful selection of saponin chemistry. In particular, HC supports magnetically aligned bicelles with high anionic lipid content, controlled access to parallel alignment via lanthanide doping, and strong resilience to Ca^2+^, providing a promising route toward more biomimetic alignment media for membrane-protein structural studies. More broadly, the distinct behaviors observed for GA, HC, and CQS define practical design rules for expanding saponin bicelles into experimental regimes where lipid composition, alignment orientation, and ionic conditions are critical constraints.

## Acknowledgements

This work was funded by the NIH (R35GM13973 to A.R.). A portion of this work was performed at the National High Magnetic Field Laboratory, which is supported by National Science Foundation Cooperative Agreement No. DMR-2128556 and the State of Florida.

## Notes

### Competing Interest Statement

The authors have declared no competing interest.

