## Supporting Information for "Saponin Chemistry Controls Anionic Lipid Tolerance and Divalent Metal Ion Responses in Magnetically Alignable Bicelles"

***Corresponding Author:**

**Supporting Information**

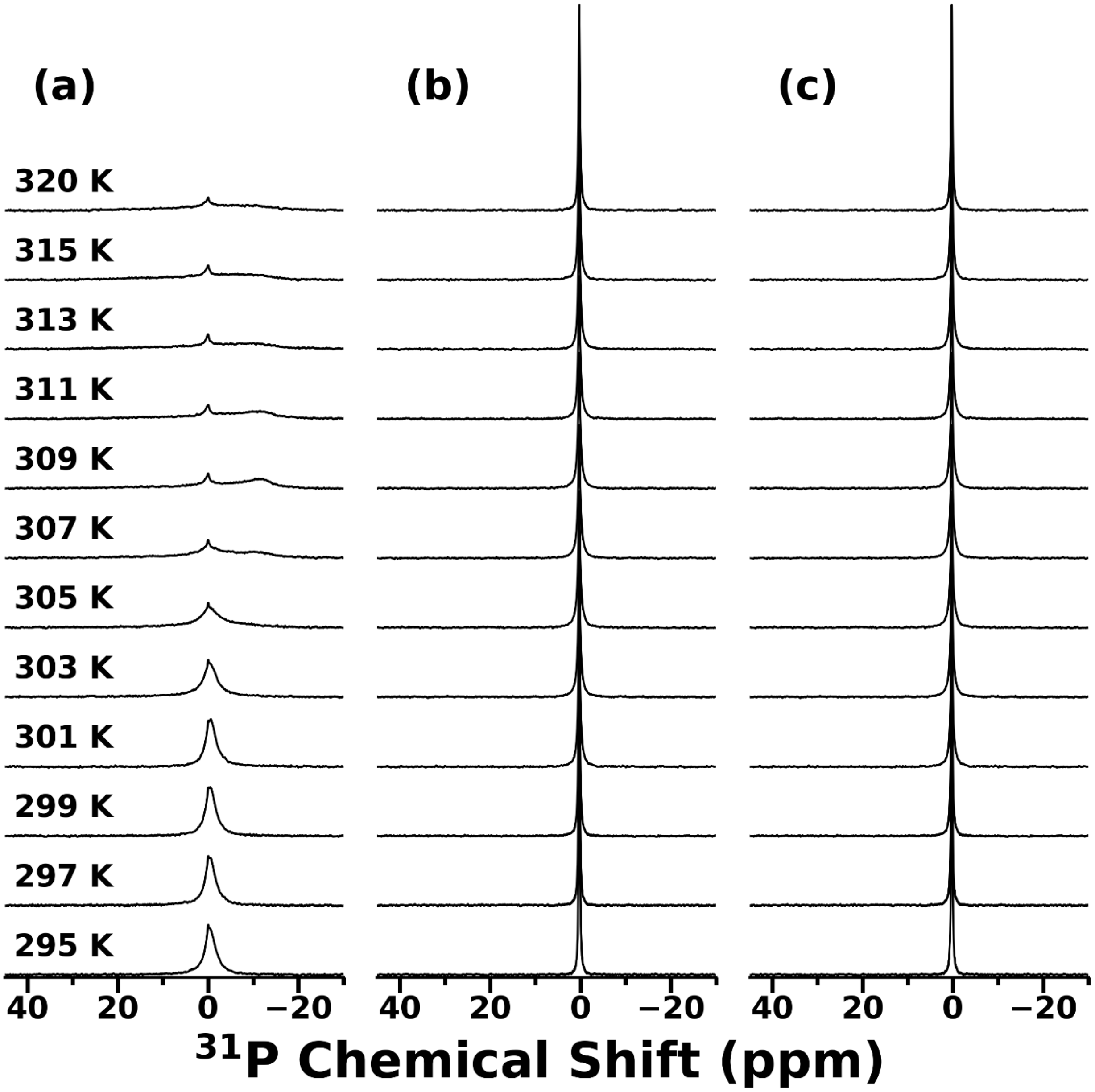


Figure S1. Temperature-dependent $^{31}$P NMR spectra of 100 mg/mL DMPG plus (a) 50 mg/mL CQS, (b) 20 mg/mL GA, or (c) 20 mg/mL HC. All spectra were collected with a 400 MHz spectrometer.

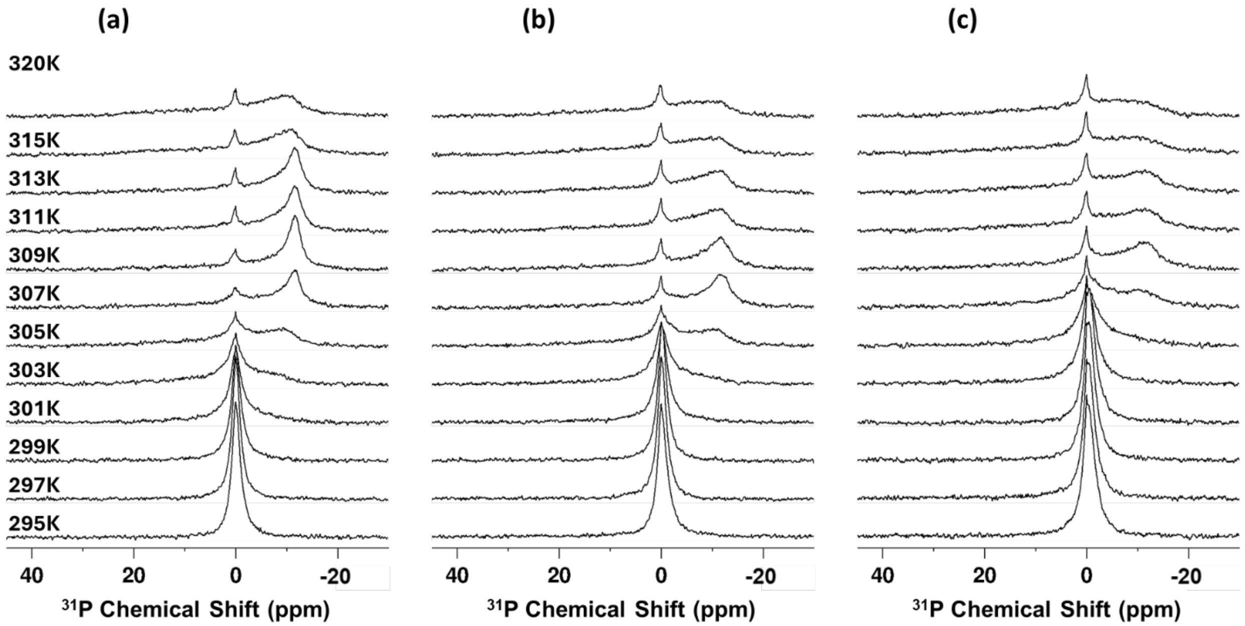


Figure S2. Effect of CQS on the temperature-dependent phase behavior of DMPG. $^{31}$P NMR spectra of mixtures of DMPG and CQS were acquired between 295 K and 320 K with a 400 MHz spectrometer. Samples were prepared with 100 mg/mL DMPG and (a) 16.7% (20 mg/mL), (b) 23.1% (30 mg/mL), or (c) 33.3% (50 mg/mL) CQS by mass in a tris buffer (10 mM tris, 100 mM NaCl, pH 7.4).

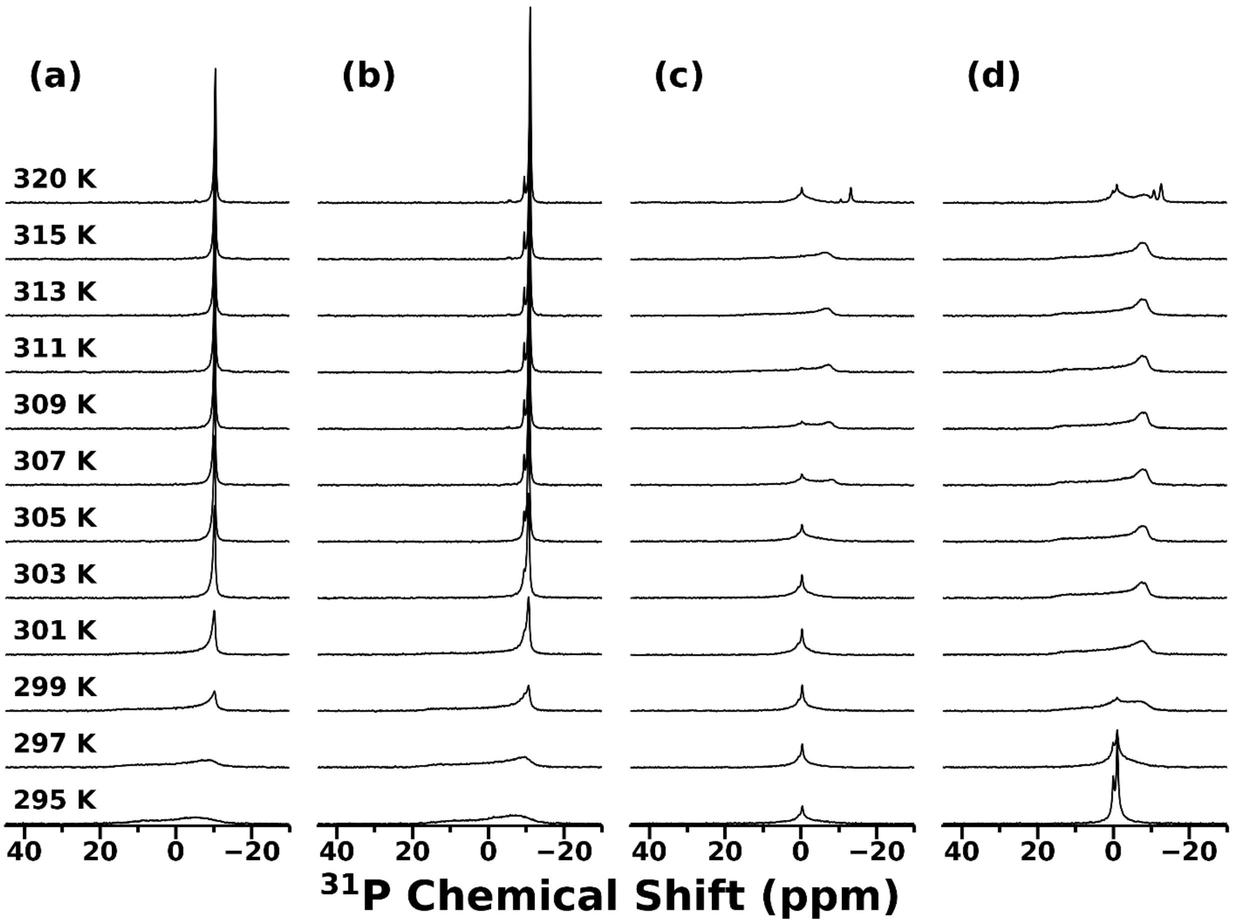


Figure S3. Effect of DMPG on bicelle formation by mixtures of DMPC and GA. Temperature-dependent $^{31}$P NMR spectra were collected with a 400 MHz spectrometer for samples containing 20 mg/mL GA and 100 mg/mL total phospholipid. The phospholipid comprised DMPC and (a) 0%, (b) 10%), (c) 15%, or (d) 30% DMPG.

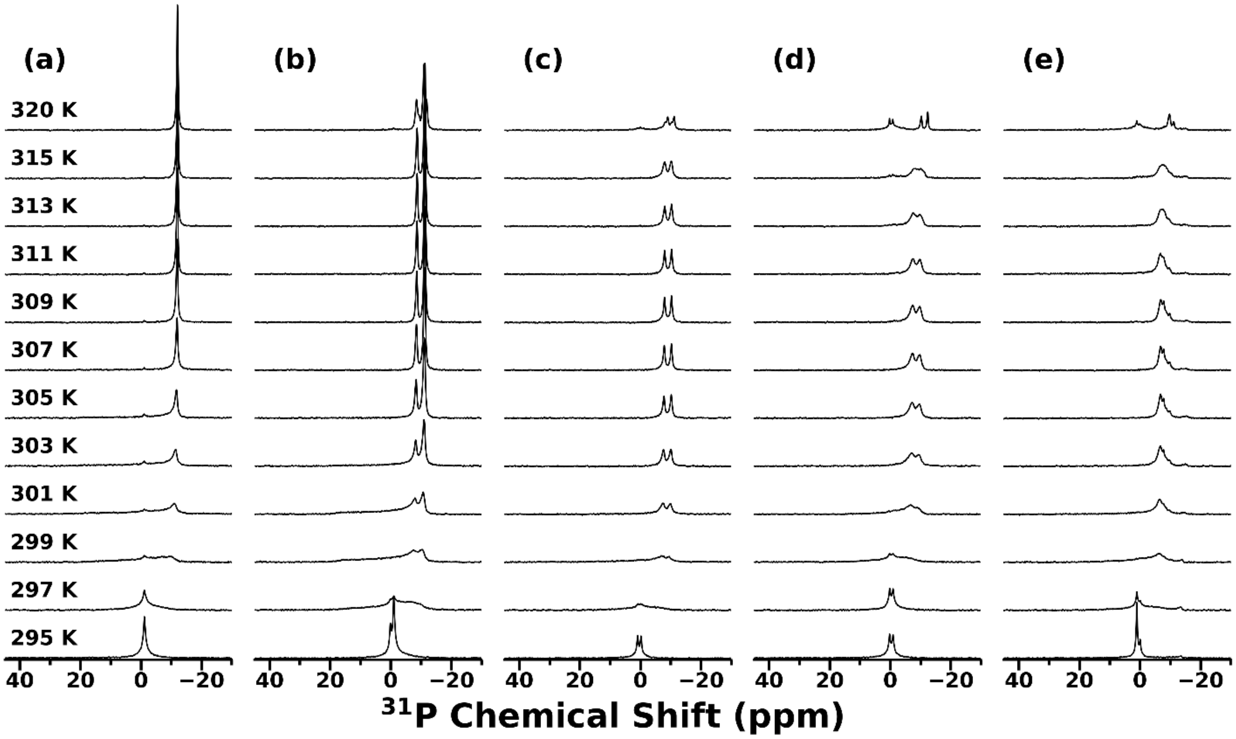


Figure S4. Effect of DMPG on bicelle formation by mixtures of DMPC and HC. Temperature-dependent $^{31}P$ NMR spectra were collected with a 400 MHz spectrometer for samples containing 20 mg/mL HC and 100 mg/mL total phospholipid. The phospholipid comprised DMPC and (a) 0%, (b) 30%), (c) 50%, (d) 55%, or (e) 70% DMPG.

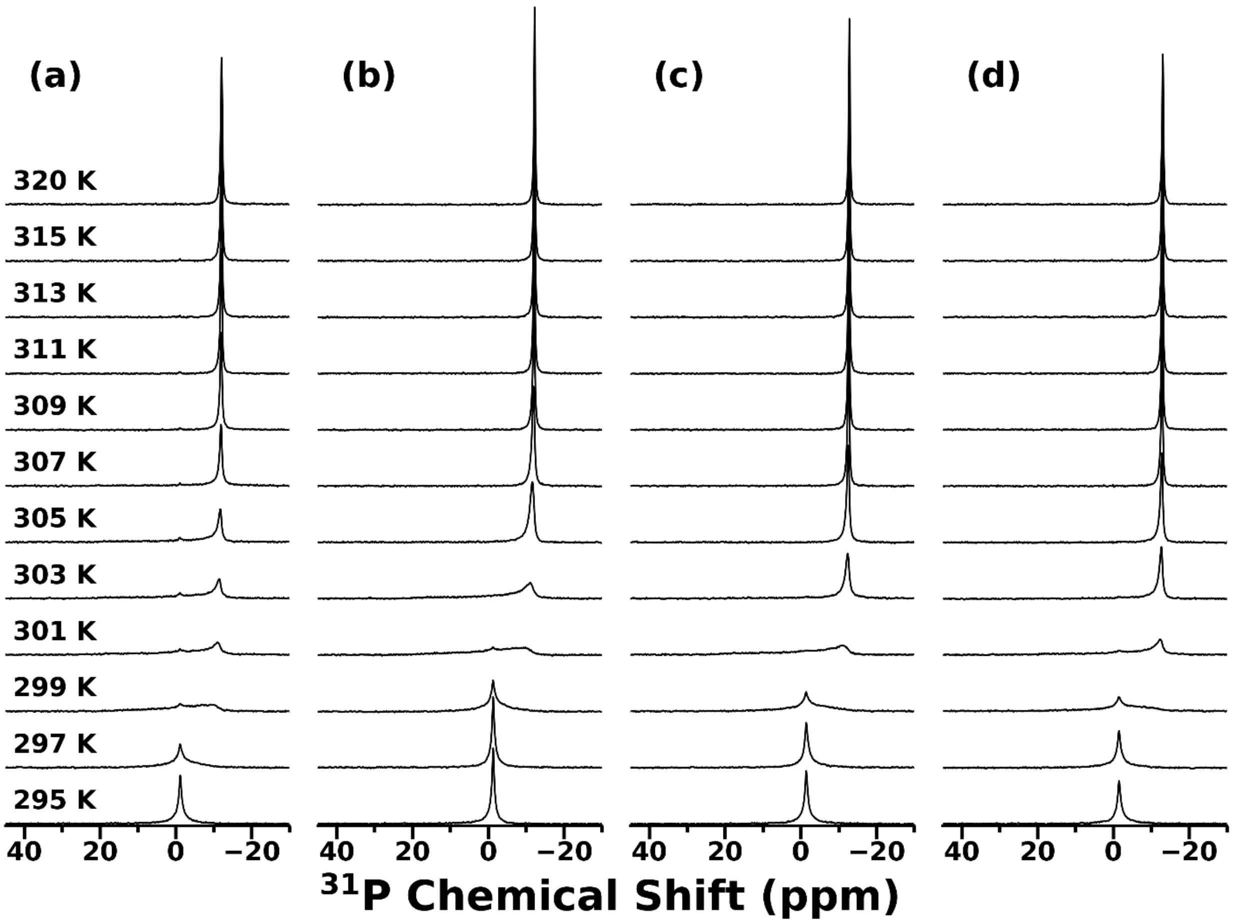


Figure S5. Effect of $Ca^{2+}$ on the stability of saponin bicelles. Temperature-dependent $^{31}P$ NMR spectra were collected for samples containing 100 mg/mL DMPC, 20 mg/mL HC, and (a) 0 mM, (b) 10 mM, (c) 50 mM, or (d) 100 mM $CaCl_2$.

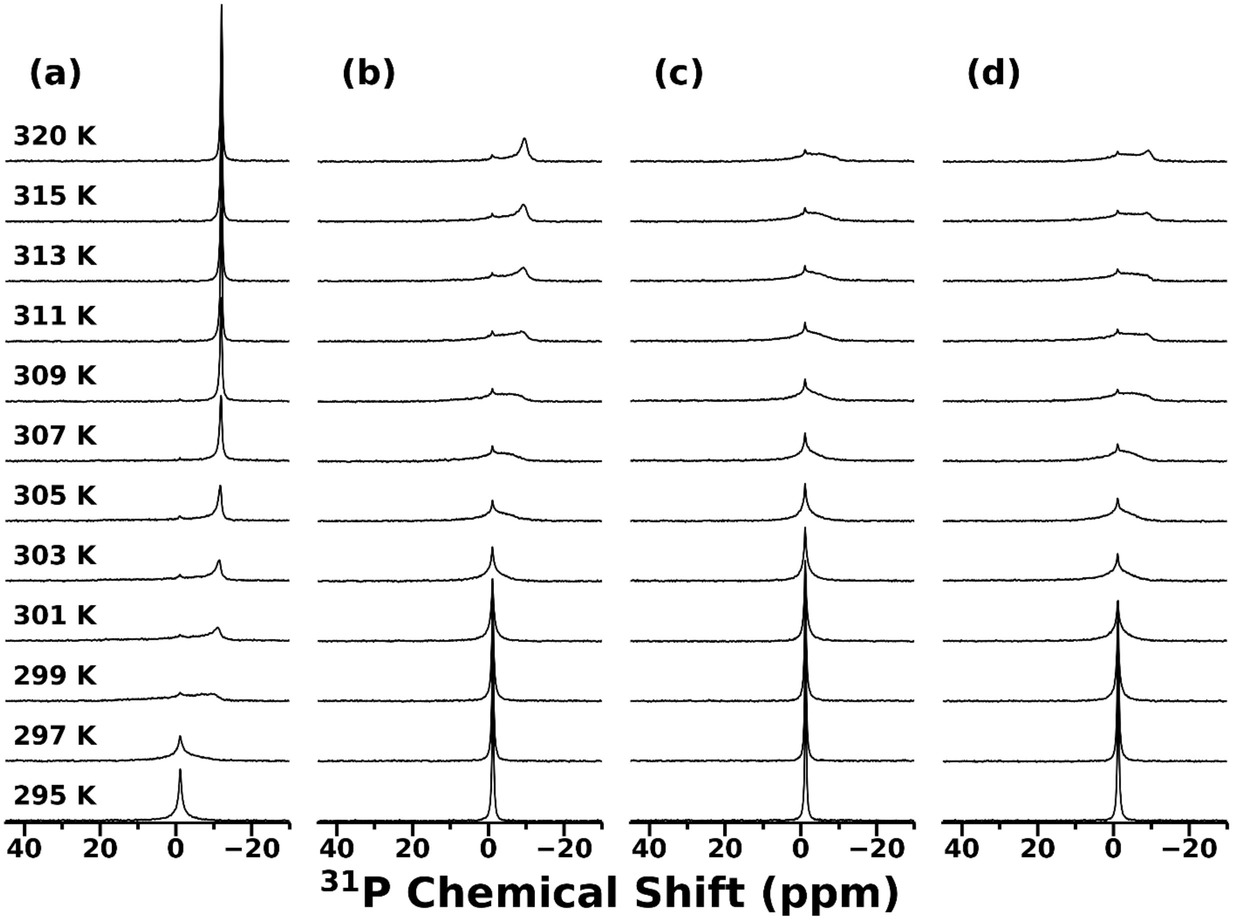


Figure S6. Effect of $Mg^{2+}$ on the stability of saponin bicelles. Temperature-dependent $^{31}P$ NMR spectra were collected for samples containing 100 mg/mL DMPC, 20 mg/mL HC, and (a) 0 mM, (b) 10 mM, (c) 50 mM, or (d) 100 mM $MgCl_2$.

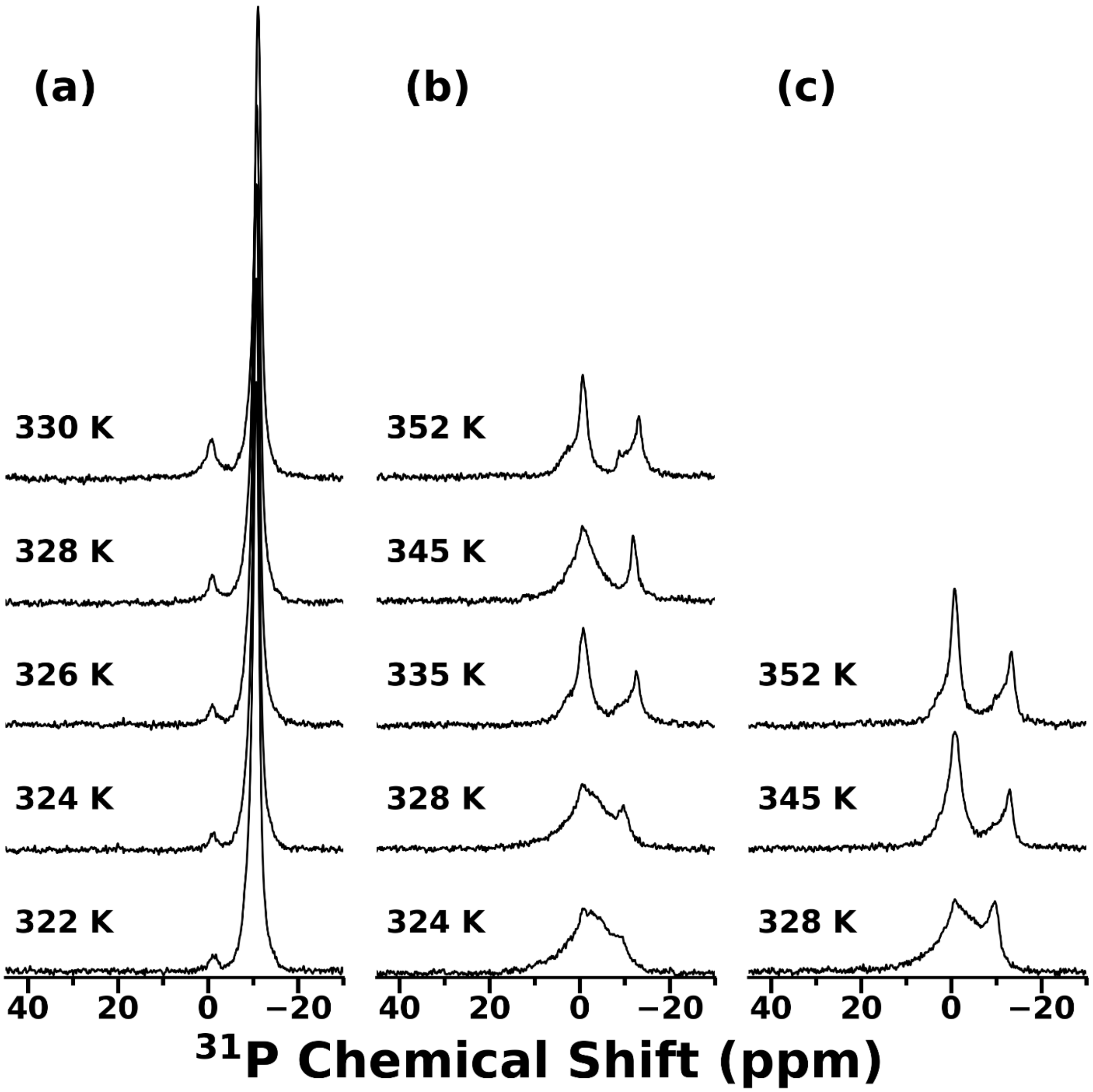


Figure S7. Effect of $Mg^{2+}$ on the stability of saponin bicelles above 320 K. Temperature-dependent $^{31}P$ NMR spectra were collected for samples containing 100 mg/mL DMPC, 20 mg/mL HC, and (a) 10 mM, (b) 50 mM, or (c) 100 mM.

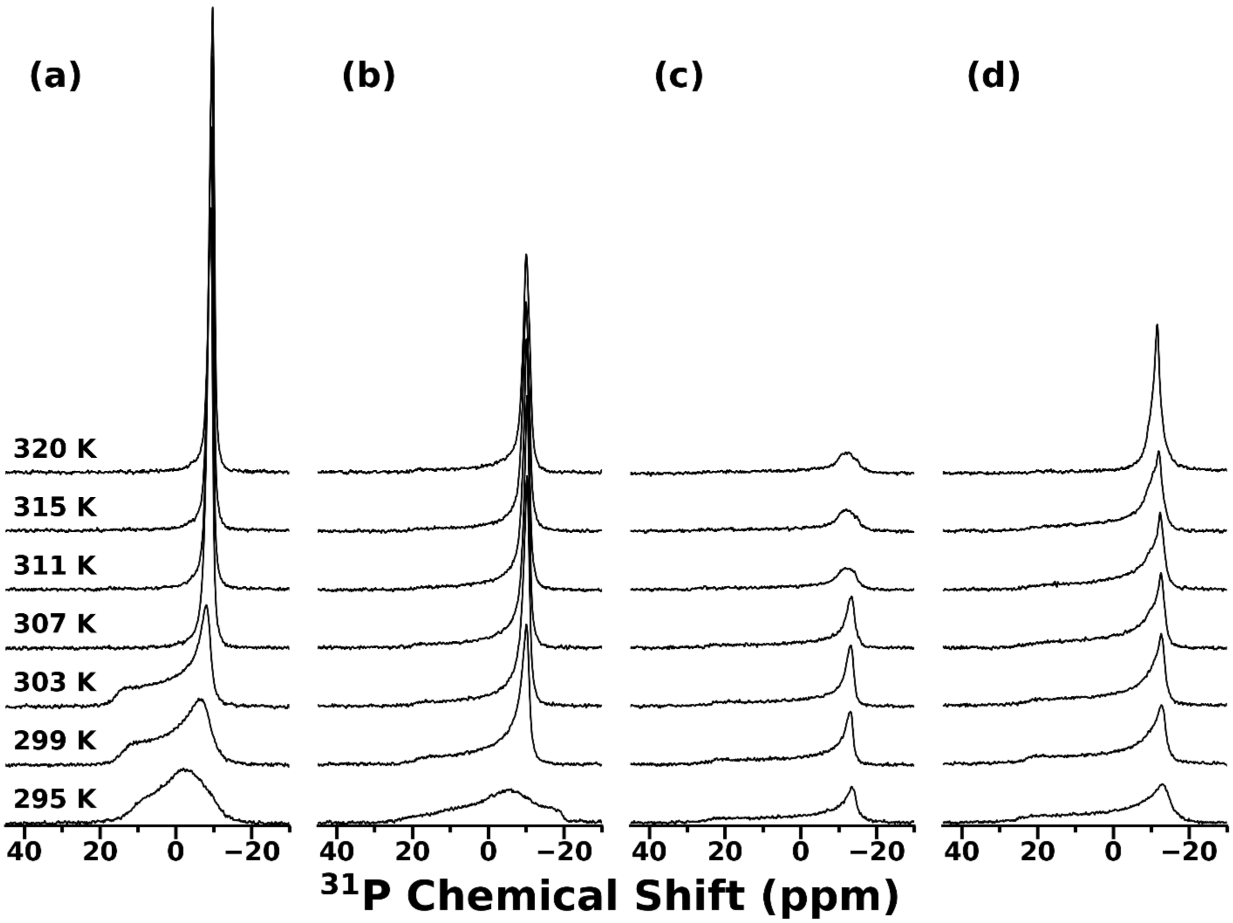

Figure S8. Effect of $Ca^{2+}$ on the stability of saponin bicelles. Temperature-dependent $^{31}P$ NMR spectra were collected for samples containing 100 mg/mL DMPC, 20 mg/mL GA, and (a) 0 mM, (b) 10 mM, (c) 50 mM, or (d) 100 mM $CaCl_2$.

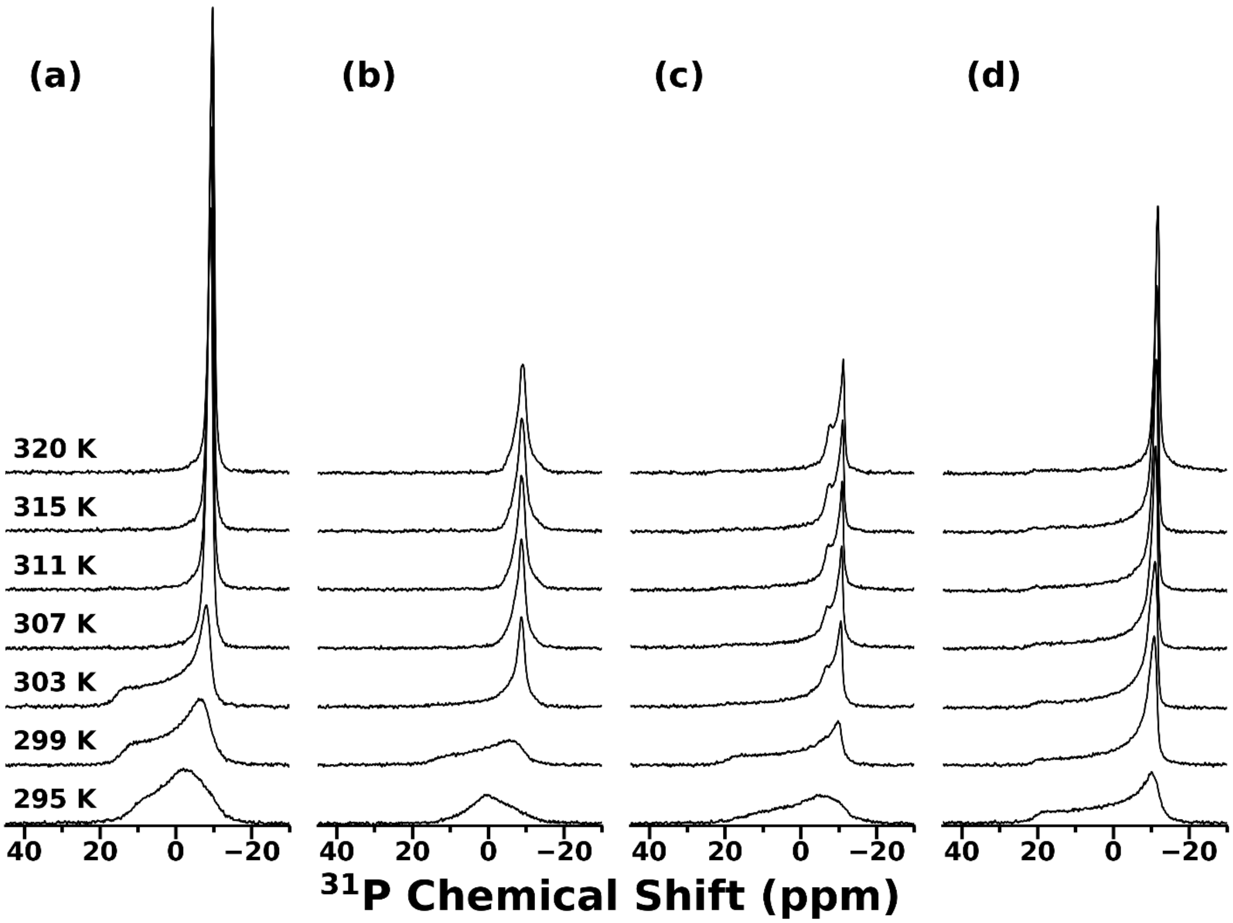


Figure S9. Effect of $Mg^{2+}$ on the stability of saponin bicelles. Temperature-dependent $^{31}P$ NMR spectra were collected for samples containing 100 mg/mL DMPC, 20 mg/mL GA, and (a) 0 mM, (b) 10 mM, (c) 50 mM, or (d) 100 mM $MgCl_2$.
